# Protein sensors of bacterial kinase activity reveal antibiotic-dependent kinase activation in single cells

**DOI:** 10.1101/2021.05.27.446034

**Authors:** Christine R. Zheng, Abhyudai Singh, Alexandra Libby, Pamela A. Silver, Elizabeth A. Libby

## Abstract

A lack of direct single-cell readouts for bacterial kinase activity remains a major barrier to our understanding of most signaling systems. At the single-cell-level, protein kinase activity is typically inferred by the activity of downstream transcriptional reporters. Complicating this approach *in vivo*, promoters are often co-regulated by several pathways, making the activity of a specific kinase difficult to deconvolve. Here, we have designed and constructed new, direct and specific sensors of bacterial kinase activity, including FRET-based sensors, as well as a synthetic transcription factor that responds to phosphorylation. We demonstrate the utility of these reporters in measuring kinase activity in population-based and single-cell assays during various growth phases and antibiotic treatments. These sensors respond to a highly conserved bacterial Ser/Thr kinase, PrkC that has no known dedicated transcription factor and whose regulon is known to be convolved with an essential signaling system. We used these new sensors to measure PrkC activity in colonies, bulk culture, and single cells. Together these new sensors provide evidence for considerable heterogeneity in PrkC activity in actively growing populations. We further demonstrate that PrkC activity increases in response to a cell-wall active antibiotic that blocks the late steps in peptidoglycan synthesis (cefotaxime), but not the early steps (fosfomycin). This is consistent with a model where PrkC senses and responds to blocks in the extracellular steps in cell wall synthesis. As the design of these phosphorylation sensors is modular, we anticipate that this work may have broad applications to other bacterial signaling systems in the future.

## Introduction

Bacteria use signaling systems to sense and respond to their environment. These sensing and response systems allow bacteria to survive in changing environments where nutrient availability and environmental stresses, such as antibiotics, are present^1,2^. When the activity of these signaling systems is disrupted, bacteria are often less adaptable to stress and more susceptible to antibiotics. Furthermore, signaling systems are also implicated in many virulence pathways of clinically important pathogens^3,4^. Therefore, understanding the regulation and activity of bacterial signaling systems is important for developing new and improved antimicrobial strategies. In this regard, the conserved PASTA containing Ser/Thr kinases in gram-positive bacteria are particularly important and understudied. They are often essential and strongly implicated in regulating virulence and antibiotic tolerance ^3,5-7^.

Currently, bacterial kinase activity can be directly measured in bulk assays, such as ^32^P labeling and phosphorylation-dependent gel shifts (Figure 1A). These methods have confirmed that phosphorylation is a widespread and important regulatory mechanism in bacteria and have been the gold standard for measuring activity both *in vitro* and *in vivo* ^2,8^. However, bulk measurements can mask individual cell variations in kinase activity in a population. Cell-to-cell variability in kinase activity likely impacts downstream regulatory pathways and subsequent phenotypes such as virulence and antibiotic tolerance^9-11^.

**Figure 1:**
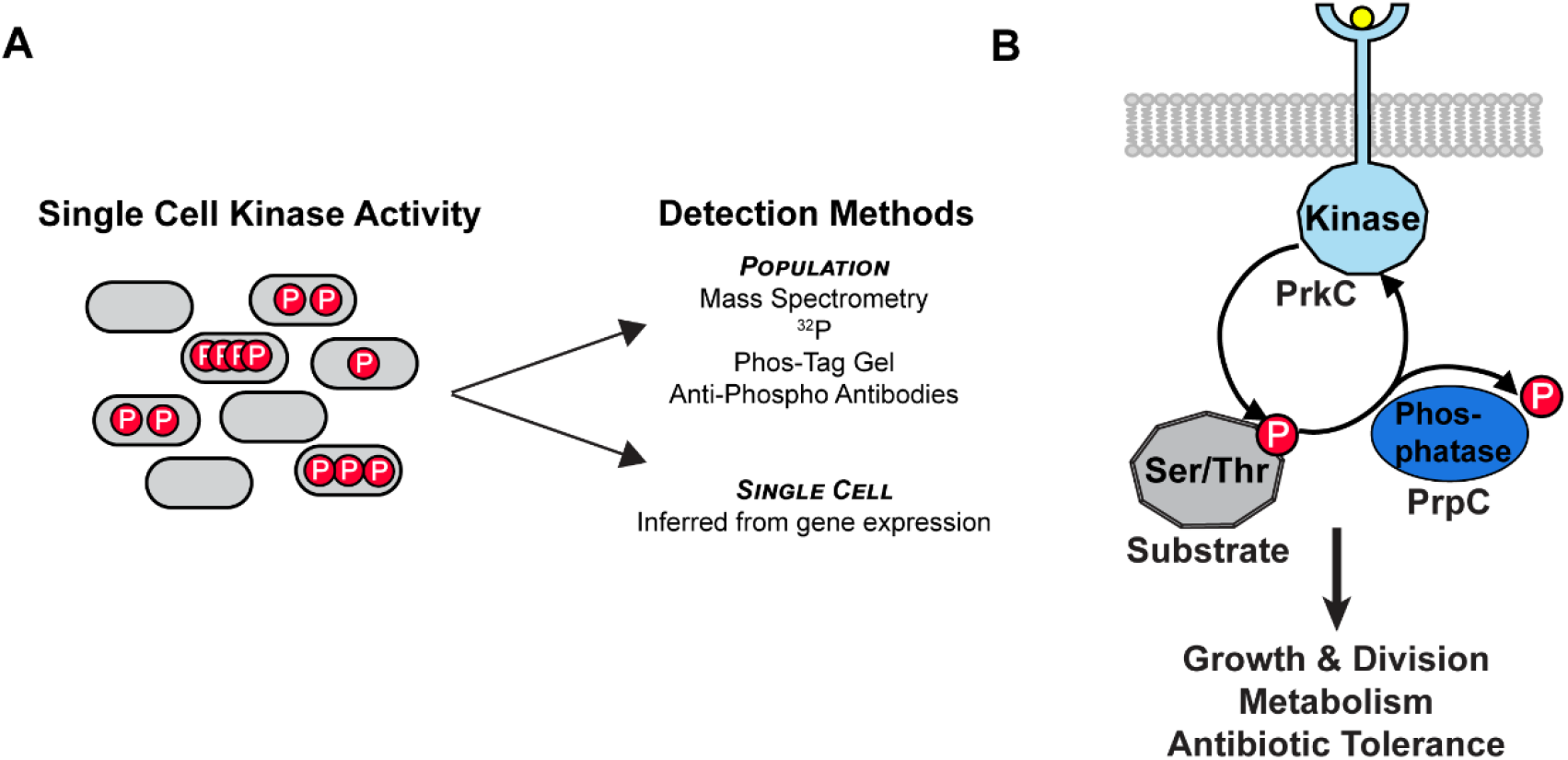
Detection of Bacterial Ser/Thr kinase activity. **A)** Established methods of measuring bacterial signaling system activity. Individual cells within a bacterial population may differ in the amount of net kinase activity, resulting in important phenotypic differences. Most established methods are population-based, such as mass spectrometry or radiolabeled ATP (^32^P). Single cell measurements of kinase activity can be inferred indirectly from gene expression of known regulated genes. **B)** Bacteria have highly conserved Ser/Thr kinase-phophatase pairs that reversibly phosphorylate target substrates. Shown is an example of a Hanks type Ser/Thr kinase pair from *B. subtilis*, PrkC/PrpC.

However, we currently lack direct measures of kinase activity in individual cells. Instead, activity is inferred from transcriptional activity of native operons that are necessarily convolved with the activity of other transcriptional regulators (Figure 1A). Several approaches have been developed to address this challenge including measuring the transcriptional response of multiple genes simultaneously, and using genetics to disentangle multiple regulatory pathways^12^. Additional less commonly used methods include quantifying transcription factor binding at specific promoters^13{Batchelor, 2006 #50^} and intermolecular FRET studies to quantify protein interactions, most notably in the *E. coli* chemotaxis pathway^14-16^. However, these methods are generally labor intensive and also can be strongly subject to intrinsic noise in cellular processes and the potentially pleiotropic effects of mutational studies.

Synthetic sensors of eukaryotic kinase activity have been implemented in some systems and have revealed spatial-temporal dynamics of various kinases, including Abl/EGFR, Aurora B, Akt, ATM, PKC, PKA, and Ras among others^17^.The general design of these sensors takes advantage of known interactions between phosphopeptides and binding domains to change the conformation of a synthetic sensor protein in a phosphorylation-dependent fashion. For example, the PKC sensor CKAR^18^ uses the interaction between phosphopeptides with a conserved motif to bind to a fork-head associated (FHA2) domain to reveal conformation dependent FRET signals. This general sensor design has been shown to be adaptable to related kinases by changing the substrate phosphopeptide to one specific for the kinase of interest (e.g., aPKC ^19^ and δPKC ^20^). This type of intramolecular FRET sensor has proven more adaptable and with generally greater dynamic range than has been achieved with intermolecular FRET studies *in vivo*.

In bacteria, Ser/Thr phosphorylation is relatively stable, compared to Asp phosphorylation (e.g. from two component signaling systems)^2^ simplifying the design of phosphorylation on-off switches. Ser/Thr phosphorylation systems often have dedicated phosphatases, providing separability between kinase and phosphatase activity. Additionally, a specific family of Ser/Thr kinases, the Hanks-type or “eukaryotic-like” (eSTK), is universally conserved across gram-positive bacteria, and are essential for growth or virulence in many clinically important pathogens ^3,7,21^. Members of this kinase family are also known to be involved in processes where phenotypic variability across populations plays an important physiological role, such as virulence and antibiotic resistance^6^. The regulation of these systems is also generally poorly understood because they often lack identified dedicated transcription factors^5^. Therefore, we sought to develop new sensors for this family of bacterial kinases to directly measure the cell-to-cell variability in Ser/Thr kinase system activity.

For our system, we chose the kinase PrkC and its partner phosphatase PrpC from *B. subtilis*^22^ (Figure 1B). PrkC is known to be involved in regulation of cell wall synthesis^23^ but its activity at the single-cell-level and role in antibiotic response has remained unclear. A major factor complicating our understanding of PrkC has been that its transcriptional regulation is largely convolved with the activity of an essential two-component system WalRK ^24^. Observed variation in gene expression in the WalR regulon gene *sasA* indicates that PrkC activity plays a key role in the antibiotic tolerance mediated by this gene ^10^. Here, we developed FRET-based sensors and new synthetic transcription factors designed to change in activity in response to phosphorylation, revealing kinase activity in colonies, bulk culture, at the single-cell-level, and during antibiotic treatment. In doing so these new sensors provide evidence that PrkC increases in activity in response to antibiotics that block the late steps in peptidoglycan synthesis.

## Results

### Design and optimization of FRET sensors for Ser/Thr kinase activity

FRET-based kinase sensors have the following key design features contained within four domains: two fluorophores capable of resonance energy transfer (FRET) when brought into close proximity, a forkhead associated domain (FHA) which binds phosphorylated proteins, and a short phosphorylatable substrate sequence^8^ (Figure 2A). Each sub-domain is connected by a flexible linker region ^18,25^. This allows the protein to change conformation upon phosphorylation. In this design, in the absence of phosphorylation the fluorophores (e.g. CFP and YFP) can be in close enough proximity to allow the emission from CFP to excite YFP, resulting in 1) a loss of CFP emission at 475 nm, and 2) an increase in YFP emission at 527 (FRET). Phosphorylation results in the substrate binding to the FHA domain, increasing the distance between the CFP and YFP, preventing the energy transfer, and resulting in a loss of FRET.

**Figure 2:**
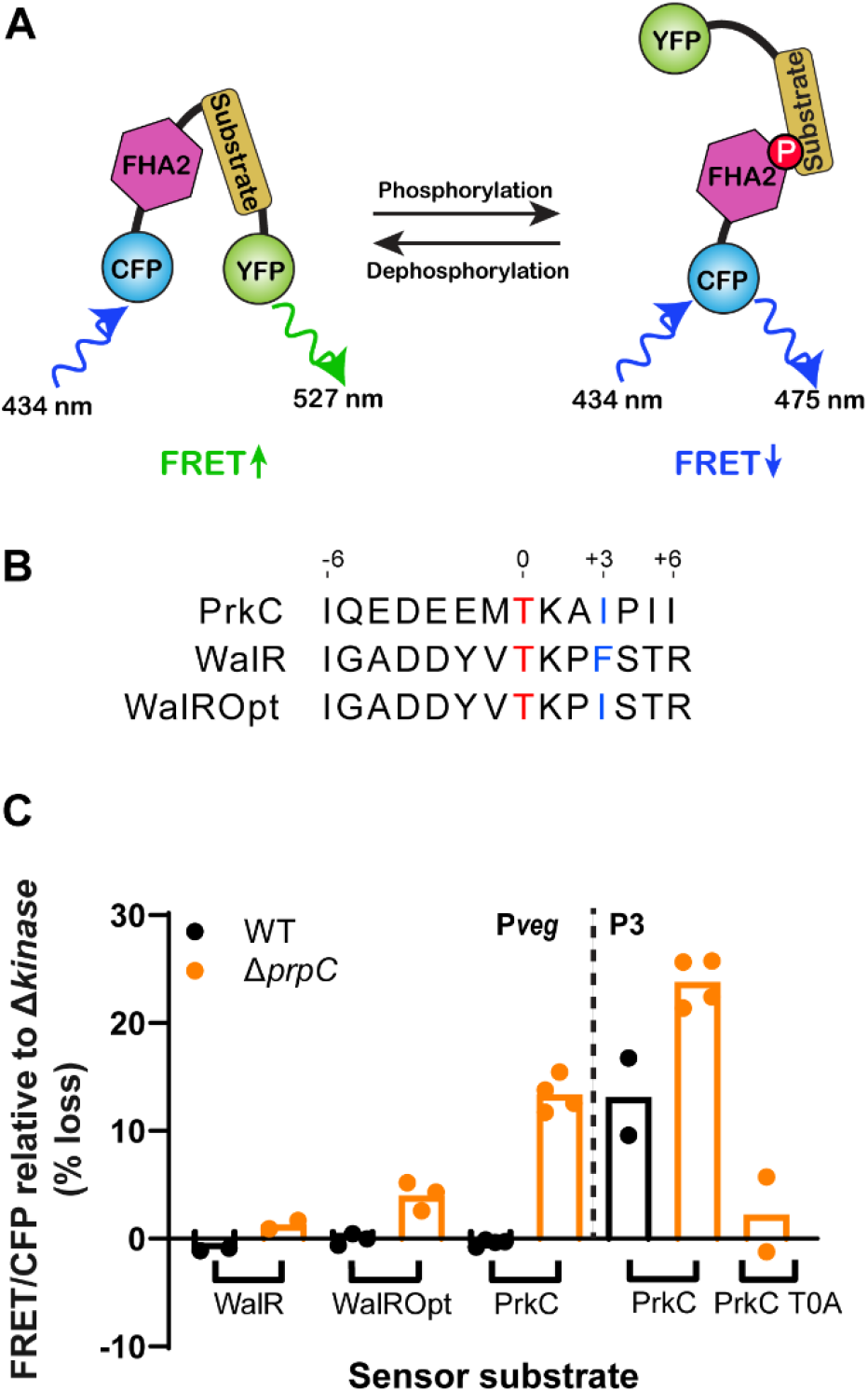
Design and implementation of FRET sensors for direct measurement of kinase activity. **A)** Schematic of an intramolecular FRET sensor responsive to kinase activity consisting of CFP and YFP fused to a phospho-binding FHA2 domain and a phosphorylation substrate for the kinase. Under conditions with low amounts of kinase activity the relative flexibility of the sensor protein allows FRET between CFP and YFP when they are in close proximity, resulting in emission from YFP at ∼527 nm when CFP is excited at ∼434 nm. Under conditions with high kinase activity, the target substrate is phosphorylated, causing it to bind the FHA2 domain, separating the CFP and YFP, resulting in a loss of FRET. **B)** Candidate sensor substrates for the Hanks type Ser/Thr kinase PrkC. Candidate substrates were chosen based on reported phosphorylation sites for PrkC itself, “PrkC” and the response regulator WalR, “WalR”. Additionally, a version of the WalR substrate with a F to I mutation in the +3 position relative to the phosphosite was chosen to increase affinity of the phosphorylated substrate to the FHA2 domain, “WalROpt”. **C)** Relative change in FRET sensor activity observed using the WalR and WalROpt substrates in a high phosphorylation state: Δphosphatase, Δ*prpC*. For each substrate the percent change in FRET/CFP signal was calculated relative to the low phosphorylation condition: a Δkinase background, Δ(*prpC-prkC*). Each dot represents the mean values of the FRET/CFP ratio from a population of ∼100 cells in an independent experiment. Right: change in FRET sensor activity with high levels of sensor expression (P_veg_) Left: change in activity for the PrkC sensor and a PrkC T0A sensor at ∼5x lower expression (P3).

The bacterial kinase PrkC from *Bacillus subtilis* is compatible with this general sensor design. Several studies identified target peptide substrates of PrkC and the associated phosphorylated residues ^26-28^. Some of these phosphorylation sites had also been verified *in vivo*, suggesting they could serve as a sensor of native kinase activity as a piece of a synthetic sensor protein. Importantly, these identified phosphorylation sites and their surrounding residues also conformed to the general features of phosphopeptides that bind FHA domains^29^. In particular, some of the best characterized sites *in vivo*, on PrkC itself, and on the response regular WalR, were compatible with FHA2 (Figure 2B). For the PrkC phosphosite in “IQEDEEM**T**KAIPII”, there is an I in the +3 position, which is believed to increase binding affinity to FHA2 domains.

The PrkC phosphosite in WalR was of particular interest, because WalR activity is currently used as a method to infer PrkC activity in vivo^24^. The peptide IGADDYVTKPFSTR did not have an optimal I in the +3 position, so we created an additional substrate peptide “WalR Opt” where the naturally occurring F in the +3 position is replaced by an I.

Using these three phosphorylatable substrates, we constructed prototype FRET sensors of PrkC activity for expression and sensing in *B. subtilis* based on the aurora B sensor^30^. In order to create a working bacterial sensor, it was necessary to codon optimize the entire protein and exchange the fluorophores for ones that work well in *B. subtilis*. For the initial construct, a strong species-specific constitutive promoter (P_*veg*_) and ribosome binding site (RBS) were also added. Once designed, the entire genetic construct was directly synthesized and cloned into a *B. subtilis* shuttle vector and subsequently integrated in single copy in the chromosome at *sacA*.

Each sensor was then tested for changes in FRET activity in high PrkC phosphorylation, Δphosphatase or Δ*prpC*, and low PrkC phosphorylation, Δphosphatase Δkinase or Δ(*prpC-prkC*), genetic backgrounds. The Δphosphatase background results in high levels of phosphorylation, as Ser/Thr phosphorylation are relatively stable and are not being actively removed by the phosphatase PrpC. The Δphosphatase Δkinase background allows measurement of the contribution to the high phosphorylation signal that is also PrkC-specific, as in the Δ(*prpC-prkC*) background PrkC is absent in addition to the phosphatase. In wild type populations (WT), both the kinase and phosphatase are present and active.

We found that of the three FRET sensors, the one with the PrkC derived substrate consistently showed the largest change in FRET between the high phosphorylation Δ(*prpC*) and low phosphorylation Δ(*prpC-prkC*) genetic backgrounds in log phase growth (Figure 2C, left, orange bars and dots). In contrast, the WalR substrate showed nearly no change. Furthermore, exchanging the F for I in the +3 position relative to the phosphosite, “WalR Opt”, produced only a modest increase in signal. Interestingly, in WT backgrounds (Figure 2C, left, black bars and dots) all 3 sensors showed no change in mean activity relative to the Δkinase background. Other work with transcriptional reporters has suggested that the average activity level in WT populations is detectable, but comparable to the Δkinase background. Taken together, these results indicate the FRET sensor is changing activity in response to PrkC, but that the sensitivity of the system needed improvement.

Based on the observation that transcriptional reporters and the FRET sensors show little to no change between Δkinase and WT populations, while the Δphosphatase populations consistently show a robust effect, we surmised that net kinase activity is low in WT populations. Therefore, our initial strategy of producing an abundance of FRET sensor proteins from a strong promoter may swamp any detectable signal with unphosphorylated sensor. We therefore chose our best performing sensor, “PrkC” together with a promoter-RBS combination “P3” that should reduce expression ∼5 fold ^31^. We then tested our system in the high phosphorylation, Δ*prpC*, and low phosphorylation, Δ(*prpC-prkC*), genetic backgrounds and found that the reduction in sensor population increased the sensitivity in both WT and Δ*prpC* populations, resulting in a measurable change in mean FRET (Figure 2C, right). To confirm the result is dependent on the T phosphosite of the sensor, we also created a version with a T to A substitution that shows a much lower average change in activity (mean ± SEM: Δ*prpC* T0A ∼2 ± 3.5%; Δ*prpC* ∼24 ± 1.1%; WT 13 ± 3.6%). Together this demonstrated that we created a FRET sensor protein that changes activity in response to PrkC.

### Design and optimization of a phosphorylation-responsive LacI: LacI∼P

We next sought to create a transcription factor that *specifically* responds to PrkC activity. To do this, we used LacI, a well characterized strong repressor of the lactose (*lac*) operon in bacteria ^32^. LacI is routinely used to provide tight, well-characterized, lactose (or IPTG) dose dependent control of the *lac* promoter in *B. subtilis*^23^ by binding to two *lacO* sites^33,34^. Our results from the FRET sensor designs demonstrated that PrkC phosphorylation can control protein conformation, suggesting that inserting a phosphorylatable substrate peptide sequence and an FHA2 domain in LacI may change the conformation of LacI in a phosphorylation-dependent manner. In this design (Figure 3A), phosphorylation would cause a decrease in LacI repressing activity, leading to downstream gene activation.

**Figure 3:**
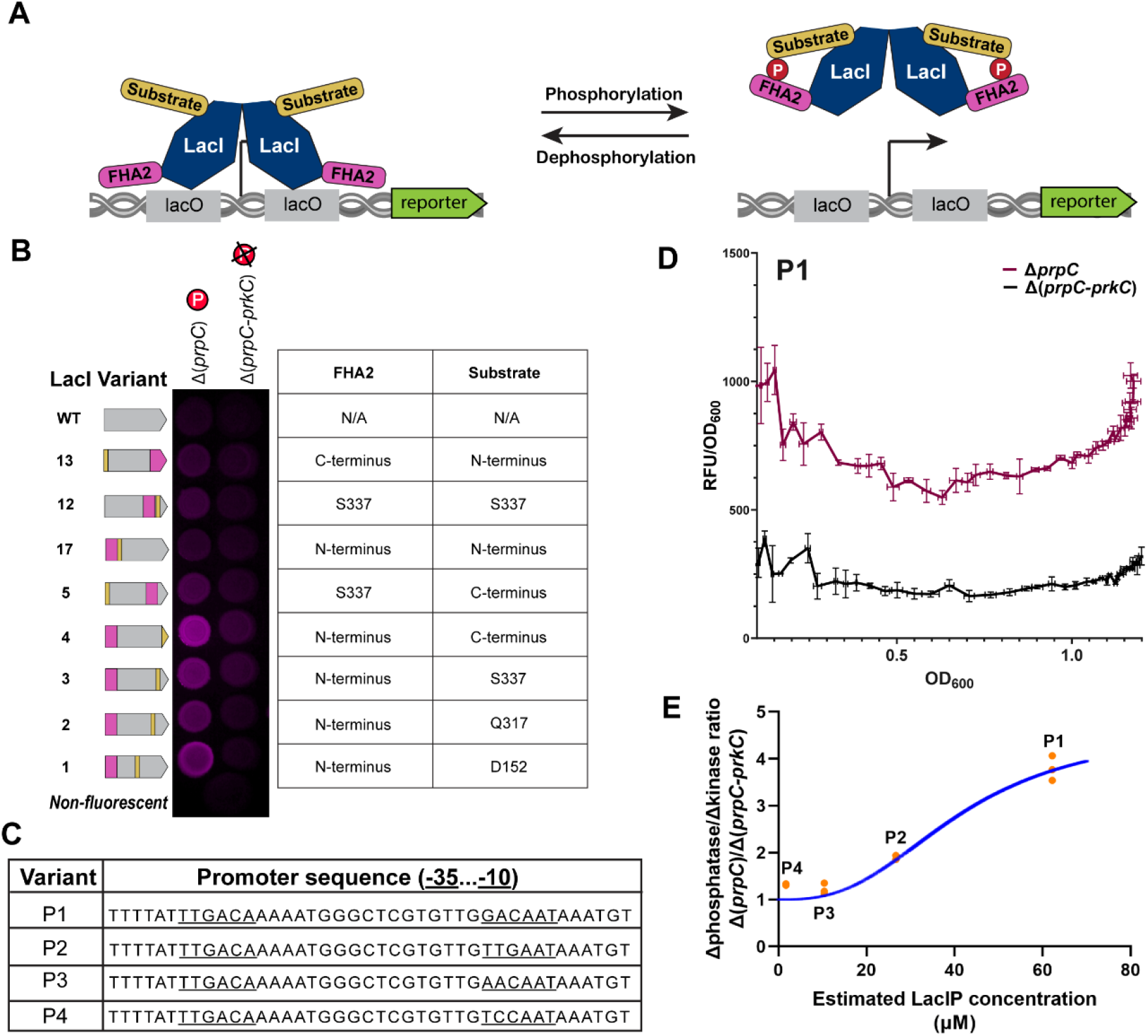
Design and testing of a synthetic phosphorylation-dependent repressor LacI∼P. **A)** Design of a phosphorylation-sensitive LacI, LacI∼P. LacI∼P was designed to respond to phosphorylation by changing conformation upon substrate region phosphorylation and subsequent binding to the FHA2 domain. The change in conformation results in a decrease in binding to the *lacO* sites and subsequent loss of repression of a downstream gene, e.g. *mcherry*. **B)** 8 design variants of LacI∼P were tested: a phosphorylatable substrate (gold) and an FHA2 domain (magenta) were inserted at various sites within LacI. The insertions are placed immediately following the amino acid listed. The DNA binding domain of LacI is near the N-terminus. For each construct, the relative repression of a downstream reporter gene (*mcherry*) was measured in high phosphorylation Δphosphatase, Δ*prpC*, and low phosphorylation Δkinase, Δ(*prpC-prkC*), backgrounds by fluorescence imaging of culture spotted on an agar plate. Wild type LacI and a non-fluorescent control are shown for reference. **C)** Promoters used to optimize measurement sensitivity of the LacI∼P to phosphorylation. Standard promoters were used with reported relative expression spanning ∼36 fold (estimated at ∼1.7 – 62.2 µM), with P1 being the highest expression, P4 the lowest. **D)** Expression of P_*lac*_-*mcherry* with P1-*lacIPv4* in Δ*prpC* and Δ(*prpC-prkC*) backgrounds as a function of OD_600_ in minimal glucose medium. Lines and bars indicate the mean and standard deviation of three biological replicates. **E)** Sensitivity of the LacI∼P (v4) system shown as the ratio of high phosphorylation/low phosphorylation state, Δ*prpC*/Δ(*prpC-prkC*), for each estimated LacI∼P expression level (P1-P4). Mean measured values for at OD_600_ ∼0.8 from 3 independent experiments similar that shown in **D** for each promoter in **C** (orange dots). We model the regulation of reporter expression by active (unphosphorylated) LacI∼P as a Hill function and use that to predict the fold change in reporter expression for high vs. low phosphorylation states (blue line). Fitting the Hill function to the data reveals a Hill coefficient of 3.

Several sites within LacI allow amino acid insertions without strongly affecting activity^35^. Using these identified sites as a guide, we inserted the phosphorylatable substrate and FHA2 domain at various positions within *lacI*^*36*^ (Figure 3B). We used a P_*lac*_*-mcherry* reporter to monitor the activity of our LacI variants. We compared the amount of mCherry in high or low phosphorylation genetic backgrounds for each permutation of insertions tested. As a control, we also verified that wild-type LacI does not respond to PrkC phosphorylation by measuring mCherry in high or low phosphorylation genetic backgrounds. Permutation #4 (FHA2 at the N-terminal DNA binding domain; substrate peptide at the C-terminus) produced the highest dynamic range between the high and low phosphorylation genetic backgrounds. We designated this construct LacI∼P.

Titration of *lacI∼P* expression revealed that we can obtain approximately a ∼4x dynamic range between the high or low phosphorylation genetic backgrounds by optimizing LacI∼P levels. We used a set of four promoter variants P1-P4 with characterized relative strengths^31^ (Figure 3C). We measured the fluorescent reporter in high Δ*prpC* or low Δ(*prpC-prkC*) phosphorylation genetic backgrounds for each promoter in a plate reader. In contrast to the established transcriptional reporters for PrkC drawn from the WalR regulon (such as P_*yocH*_), we found that the fluorescence signal is relatively stable as a function of OD_600_ (Figure 3D). We also found that varying the relative estimated LacI∼P level by changing the promoter strength led to large changes in the dynamic range of the reporter (Figure 3E, orange dots). The system with the highest level of LacI∼P, P1-*lacIP*, showed the largest dynamic range ∼4x. We compared the estimated expression level of LacI∼P and the observed dynamic range from each experiment and found that it could be fit to a Hill function with a coefficient of ∼3 (Figure 3E, blue line), demonstrating that LacI∼P exhibits cooperative behavior similar to wild type LacI^37^.

### LacIP reveals PrkC activity in colonies and single cells

In order to increase the reliability of our LacI∼P system across growth conditions, we normalized our P1-*lacIP* P_*lac*_-*yfp* reporter system to a constitutive P_*veg*_-*cfp* reporter. The addition of a constitutive reporter allows for normalization of the signal of interest (YFP) against a general reporter that corrects for general fluctuations of cellular transcription and translation (CFP), a strategy that is particularly useful when measuring variability in populations^38^. By streaking this combined system on agar plates, we were able to easily visualize differences in colony fluorescence in a high phosphorylation genetic background (Δ*prpC*, Δphosphatase) vs. a low one (Δ(*prpC-prkC*), Δkinase) (Figure 4A, upper left quadrant vs. lower left). Consistent with previous results using WalR regulon transcriptional reporters or phos-tag gels^24^, as well as our FRET sensor results in this work (Figure 2), wild type colonies show similar levels of phosphorylation to Δkinase (Figure 4A, upper right quadrant).

**Figure 4:**
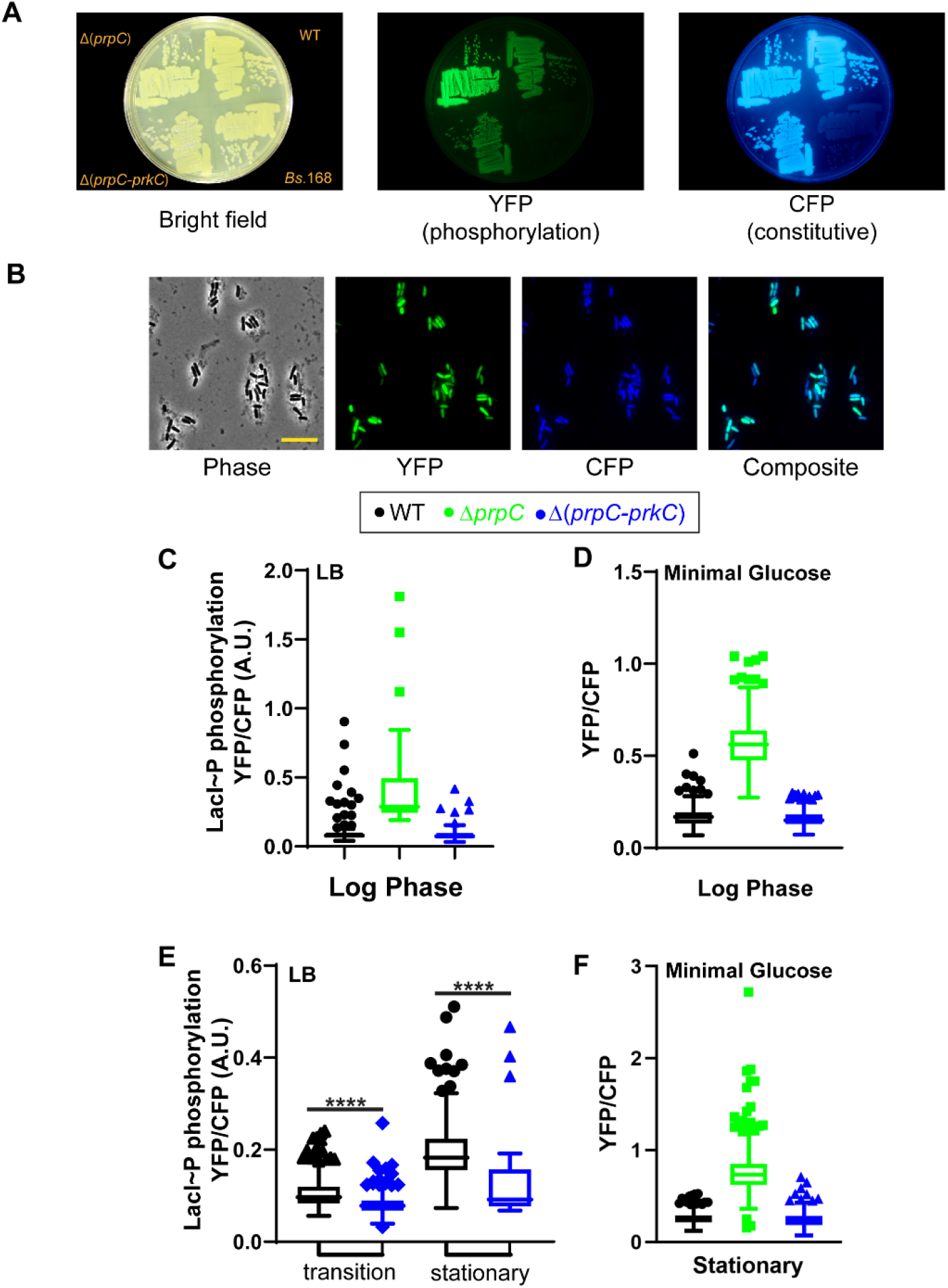
The LacI∼P system reveals growth phase dependent heterogeneity in PrkC kinase activity. **A)** System for normalizing LacI∼P signal: P_*lac*_ -*yfp* expressed in combination with P_*veg*_ -*cfp*, allowing for normalization of cell-to-cell variability in global protein expression. Strains were constructed with the promoter optimized P1 expression system from Figure 3, in otherwise wild type (WT), Δphosphatase, Δ*prpC*, and Δkinase, Δ(*prpC-prkC*) backgrounds. Representative fluorescence images of the resulting strain streaked on LB agar is shown. Left: Brightfield, Middle: YFP, Right: CFP. **B) Left:** Single cell images of the LacI∼P system from **A** in a wild type population grown to stationary phase in rich media (LB). Scale bar ∼ 10 µm. **Right:** A composite overlay of YFP (P_*lac*_ -*yfp*) and CFP (P_*veg*_ -*cfp*). **C-F)** Normalized LacI∼P activity as a function of growth media and growth phase in single cells. Shown are the distributions of the YFP/CFP ratio in single cells in a representative experiment for each genotype and growth condition. Box plots shown indicate the interquartile range and median of the data sets, whiskers indicate the Tukey interquartile range and associated outliers are also shown. In **D** the Δ*prpC* background is omitted in order to visualize the changes in WT populations. OD_600_ in LB: log ∼0.7, transition ∼3.0, stationary (varies by genotype – WT∼0.4, Δ(*prpC-prkC*) ∼0.1. OD_600_ in minimal glucose: log ∼0.5, stationary ∼3. (****, p-value <0.0001, KS-test)

Previously, transcriptional reporters used to measure PrkC activity suggested that PrkC activity is strongly media and growth phase dependent. However, the reporters used (e.g., P_*yocH*_) report on WalR activity under dual phosphorylation from both its cognate essential histidine kinase WalK and PrkC/PrpC. For example, in log phase in rich media (Luria broth) P_*yocH*_ does not show a difference in reporter activity between high and low PrkC-phosphorylation states^24^ but it does in minimal glucose media^10^. Additionally, wild type populations show relatively low average levels of net PrkC activity. This may be due to relatively uniform low levels of activity and/or masking significant cell-to-cell variability. We therefore used our LacI∼P system to quantify the distribution of PrkC activity in single cells. Visual inspection of cells grown to early stationary phase showed evidence of significant cell-to-cell variability in PrkC activity, with variability observed in the LacI∼P regulated YFP channel, that was not observed in the constitutive CFP channel (Figure 4B).

By comparing PrkC activity in both rich (LB) and minimal glucose media, we found consistent and strong differences in PrkC activity between high and low phosphorylation genetic backgrounds in log phase growth (Figure 4C and D, green and blue), indicating PrkC-dependent activity regardless of growth media. Wild type populations in both growth media show comparable levels of activity to Δkinase (LB YFP/CFP mean ± SEM: Δ*prpC* ∼0.42 ± 0.05; Δ(*prpC-prkC*) ∼0.10 ± 0.01; WT 0.11 ± 0.01 and Minimal glucose YFP/CFP mean ± SEM: Δ*prpC* ∼0.57 ± 0.01; Δ(*prpC-prkC*)∼0.16 ± 0.003; WT 0.17 ± 0.003), with rare cells showing distinctly elevated levels of phosphorylation signal. In LB however, wild type populations begin to accumulate increasing numbers of cells with elevated YFP/CFP in transition phase OD_600_∼3.0 and in stationary phase after the onset of lysis, OD_600_∼0.4 (Figure 4E). In contrast, in minimal media, the populations remain relatively similar through stationary phase (Figure 4F). This suggests that the previously reported WalR regulon transcriptional reporters were reflective of PrkC activity in minimal glucose media, but masked activity in log phase in LB^10,24,39^.

### LacIP reveals antibiotic-dependent PrkC activation

How PrkC responds to antibiotic treatment *in vivo* remains unclear. We therefore sought to determine if our LacI∼P PrkC activity reporter system could measure changes in activity under treatment with antibiotics that block early vs. late steps in peptidoglycan synthesis. Fosfomycin, a MurAA inhibitor, blocks the early, or cytoplasmic steps in the pathway^40^. Cefotaxime, a β-lactam, blocks the late steps in the pathway^41^. Fosfomycin treatment is known to deplete lipid II, whereas treatment with β-lactams such as cephalosporins block the incorporation of lipid II into the peptidoglycan sacculus^42^. Previous work has suggested that PrkC may be important for sensitivity to the 2^nd^ generation cephalosporin cefuroxime in *B. subtilis*, although modestly so^43^. Here we found PrkC plays a strong role in cefotaxime (a 3^rd^ generation cephalosporin) sensitivity in *B. subtilis*, with the low phosphorylation genetic background Δ(*prpC-prkC*) being ∼20x more sensitive than a high phosphorylation Δ*prpC* background in a test strip sensitivity assay (Figure 5A). This result is consistent with PrkC activity increasing in response to cefotaxime and subsequently exerting a protective effect.

**Figure 5:**
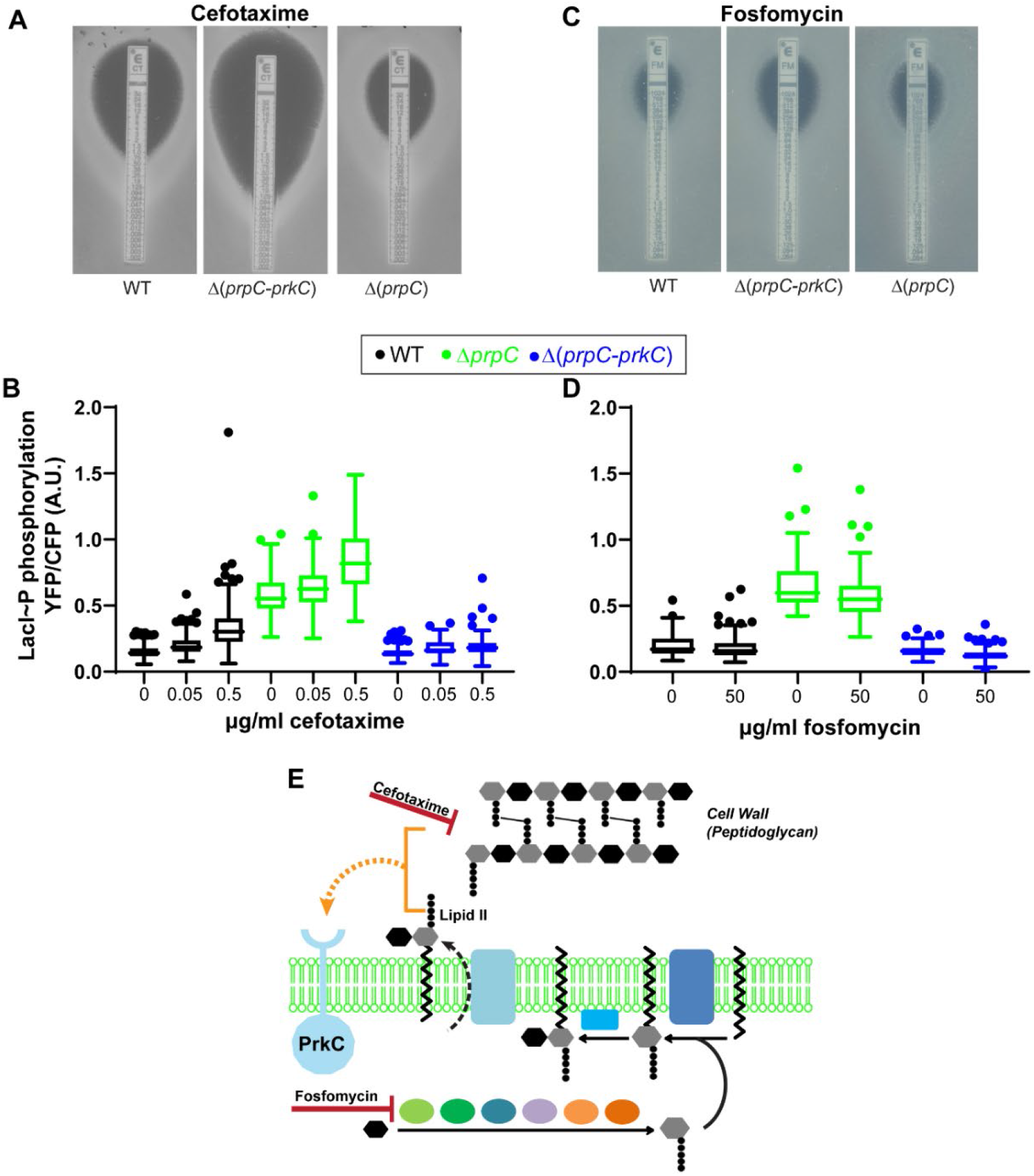
LacI∼P measures PrkC kinase activation during treatment with the β-lactam antibiotic cefotaxime. **A)** Antibiotic test strips measuring the sensitivity to the β-lactam antibiotic cefotaxime in wild type (WT), Δphosphatase, Δ*prpC*, and Δkinase, Δ(*prpC-prkC*) backgrounds on minimal glucose agar plates. Intersection of bacterial growth with the strip indicates ∼sensitivity in µg/ml. Cefotaxime inhibits the late, extracellular, steps in peptidoglycan synthesis. **B)** LacI∼P phosphorylation was inferred from normalized P_*lac*_ -*yfp* expression under treatment with increasing concentrations of cefotaxime, up to 0.5x WT MIC, in WT, Δ*prpC*, and Δ(*prpC-prkC*) backgrounds. Shown are the distributions of YFP/CFP in single cells in a representative experiment. Box plots shown indicate the interquartile range and median of the data sets, Tukey whiskers indicate the interquartile range and outliers are also shown. Each distribution is comprised of at least 125 cells, apart from Δ(*prpC-prkC*) treated populations where the antibiotic was acutely toxic (∼50 cells each). **C)** Similar to **A**, but for the antibiotic fosfomycin. Fosfomycin inhibits the early, cytoplasmic, steps in peptidoglycan synthesis. **D)** Similar to **B**, for the antibiotic fosfomycin. **E)** Model for PrkC activation. Fosfomcyin inhibits MurAA, an enzyme in the early, cytoplasmic steps of peptidoglycan synthesis, depleting lipid II. In contrast, cefotaxime blocks the late steps in peptidoglycan synthesis. PrkC may sense the availability of lipid II or another peptidoglycan precursor that accumulates during cefotaxime treatment through interactions between its extracellular receptor and the m-DAP in the stem peptide.

We therefore measured changes in PrkC activity during cefotaxime treatment using the LacI∼P system. We measured LacI∼P normalized system activity during treatments with 0, 0.05, and 0.5 µg/ml cefotaxime in minimal glucose media (Figure 5B); 0.5 µg/ml is roughly half the MIC (Figure 5A) but is well above the MIC for Δ(*prpC-prkC*). For each genotype, we measured the YFP/CFP ratio in individual cells. We found that cefotaxime produced an increase in LacI∼P reporter activity in both WT and Δ*prpC* (Δphosphatase) populations, and a much smaller effect on Δ(*prpC-prkC*) (Δkinase) populations. This is consistent with cefotaxime treatment causing an increase PrkC activity. To quantify the increase in kinase activity during cefotaxime treatment, we used data from the Δphosphatase background in our model for LacI∼P phosphorylation (Materials and methods, and Figure 3E). We estimate that the change in reporter activity at 0.5 ug/ml cefotaxime is consistent with ∼ 63-75% increase in kinase activity (95% confidence interval).

In contrast to cefotaxime, PrkC does not have a strong effect on fosfomycin sensitivity (Figure 5C). Since fosfomycin blocks the production of lipid II, a proposed activating ligand for PrkC, and the basal level of activity for PrkC is very low on average - PrkC activity may not change upon fosfomycin treatment. We used the LacI∼P system to measure PrkC activity during treatment with increasing concentrations of fosfomycin, up to ∼0.5x MIC in minimal glucose media (Figure 5D). We found that fosfomycin did not strongly change the YFP/CFP ratio, even though the cell morphology showed strong evidence of antibiotic toxicity.

## Discussion

A lack of direct single cell readouts for kinase activity remains a major barrier to our understanding of most bacterial signaling systems. In this study we focused on developing sensors for Ser/Thr kinase activity for a conserved class of signaling systems. These systems were deemed particularly tractable due the relative stability of the Ser/Thr phosphorylation and prior success in related eukaryotic systems^44^. Another key factor was the genetic separability of kinase and phosphatase activity, allowing us to easily measure the specific activity of these sensors.

Our initial FRET sensor design adapted prior work on eukaryotic systems to the kinase-phosphatase pair PrkC/PrpC in *B. subtilis* (Figure 2A). Successful adaptation required the use of a specific phosphorylatable substrate compatible with the FHA2 binding and titration of sensor expression to increase measurement sensitivity (Figure 2C). Our results indicate that our sensor is specific for PrkC, as deletion of the kinase or the partner phosphatase PrpC give opposing regulation. The most robust substrate phosphopeptide was chosen from PrkC itself, although a WalR substrate which contains the Thr101 phosphosite did yield modest activity when changed to include features that increase binding to FHA2. Together, these results indicate that in order to achieve sensitivity and specificity in bacteria, several factors need to be simultaneously optimized.

Using a conceptually similar strategy, we designed a phosphorylation-dependent repressor, LacI∼P, by inserting the phosphorylatable substrate and FHA2 in LacI (Figure 3). By choosing a specific substrate peptide for PrkC, we created a dedicated transcription factor for PrkC. This disentangled measurements of PrkC from the two-component system WalRK, whose regulon have been used to infer PrkC activity. The response regulator WalR is phosphorylated on Asp 53 by WalK and on Thr 101 by PrkC, increasing WalR activity as both an activator and repressor^24^. However, WalK-dependent phosphorylation of WalR is essential^45,46^, preventing full separation of PrkC and WalK activity by this downstream measure. A further complication is that genes in the WalR regulon are co-regulated by other transcriptional regulators, including the PhoPR TCS^47^, Spo0A^48^ and AbrB^49^. Therefore, creating a dedicated transcription factor for PrkC, allowed us to isolate and measure the activity of PrkC. This was particularly advantageous in situations where other regulators may come into play, such as changing media, growth phase, and during antibiotic treatment. Furthermore, the cooperativity of LacI-P (like LacI) can increase the sensitivity and response to phosphorylation.

Several studies had indicated that the extracellular receptor PASTA domain of PrkC and its close homologs binds to muropeptides ^50-52^, leading to the hypothesis that PrkC may monitor cell wall growth *in vivo* by responding to a peptidoglycan intermediate. To date, most *in vivo* evidence of changes in PrkC activity was obtained through anti-phosphothreoine antibodies ^23,53^, phos-tag^24^, phosphoproteomics ^27^, or downstream transcriptional changes convolved with another signaling system ^10,24^. Although changes in PrkC activity could be detected using these methods, the activity in wild-type populations was generally very low and close to that of Δkinase populations. These results could be consistent with PrkC having extremely transient, or very low levels of activity during active growth, or cell-to-cell variability masked by bulk measurements.

Here we use our specific sensors of PrkC activity to measure kinase activity in single cells as a function of growth media as well as during antibiotic treatment. In doing so, we resolve the apparent contradiction that PrkC is only active during log phase growth in minimal media, but not in rich media (LB)^10,24^ that was inferred from transcriptional activity of WalR operon genes. Here, we found using the LacI∼P reporter system that PrkC is active in both types of growth media, as well as on agar plates (Figure 4). This result is consistent with the hypothesis that PrkC senses a cell wall precursor, such as lipid II, which is expected to be present during log phase growth. Our results are also consistent with PrkC exhibiting transient bursts of activity, since the Δ*prpC* background shows consistently high levels of activity, compared to wild type backgrounds with active phosphatase activity. We also found that the LacI∼P reporter system measures changes in PrkC activity that are broadly consistent with antibiotic sensitivity of kinase and phosphatase genotypes (Figure 5). For example, a Δ(*prpC-prkC*) mutant is more sensitive to cefotaxime than a Δ*prpC*, suggesting PrkC activity may increase during cefotaxime treatment, whereas Fosfomycin does not exhibit a clear different in either antibiotic sensitivity or PrkC activity. The lack of response to fosfomycin treatment, which blocks the early steps in the pathway further supports this model and indicates that the measured increase in LacI∼P reporter system activity is not a generalized global cell wall stress response.

Since close homologs of PrkC are essential for growth and virulence in many clinically important pathogens, developing sensors of PrkC demonstrates a new modular method to probe the activity of an important class of bacterial signaling systems. Importantly, our development of a modular synthetic transcription factor overcomes hurdles present in other technically challenging approaches like FRET and therefore provides a new methodology to measure kinase signaling in single cells.

## Materials and Methods

### Strain construction

Details of strains, plasmids, and oligos used are in Table S1-S3. Plasmids were constructed via standard molecular biology techniques, including Golden Gate cloning, or by direct synthesis from IDT. We selected for successful transformants on LB containing 100 μg/mL ampicillin.

*Bacillus subtilis* strains were constructed by transformation of DNA into *B. subtilis* 168 *trpC2* and its derivatives using standard techniques^54^. Where appropriate genomic DNA was obtained using the Wizard Genomic DNA Purification Kit (Promega). We selected for successful transformants on LB Lennox containing appropriate antibiotic (10 μg/mL kanamycin, 100 μg/mL spectinomycin, 5 μg/mL chloramphenicol, or MLS).

### Media and culture conditions

*E. coli* and *B. subtilis* growth and media conditions were similar to those described previously^10,24^. Briefly, *E. coli* were grown on LB agar with an antibiotic as needed at 37°C overnight before being inoculated into LB or 2XYT liquid medium with an antibiotic as needed and grown at 37°C.

*B. subtilis* cultures were inoculated from single colonies and were grown in LB Lennox and S7 liquid media as indicated at 37°C. For the preparation of S7 growth media, we modified the Teknova MOPS Minimal Media kit (M2106) to the appropriate concentrations and supplemented with 0.1% w/v glutamic acid and 40 μg/mL L-tryptophan.

For FRET experiments, single colonies grown on LB Lennox agar overnight were inoculated into S7 media and grown in a roller drum to early log phase. For imaging purposes, the cultures were then concentrated via centrifugation (3000 rcf, 2 min) and resuspended in the original supernatant.

For antibiotic treatment experiments, 3ml cultures were grown to midlog (OD_600_∼0.4) in S7 media before being split and treated with the antibiotic at the concentration indicated for 1.5 h. Antibiotic sensitivity experiments were performed using ETest® strips (Biomerieux) on S7 agar plates at 37°C overnight.

### Microscopy and image analysis

Microscopy was performed on live cells immobilized on 1% agarose pads prepared with S7 media. Images were taken on a Nikon TE2000 or a Nikon Eclipse Ti2 microscope using a Phase contrast objective (CFI Plan Apo Lambda DM ×100 Oil, NA 1.45), an X-Cite light source, a Hamamatsu Orca ER-AG, and the following filter cubes: FRET (ex436/20, dm455lp, em535/30), CFP (ex436/20, dm455lp, em480/40), YFP (ex500/20, dm515, em535/30), and mCherry (ex560/40, dm585, em630/75).

Images were analyzed as described previously^10^ using Fiji^55^ and MATLAB to obtain cell averages for fluorescence values. Briefly, to calculate fluorescence per cell, first the pixels (or mask) of individual cells were identified. To do this, a binary image was created by thresholding the phase image to include each cell’s interior and edge. Next, individual cells were identified by discovering connected components within the binary image using MATLAB 2019b (Mathworks) function bwconncomp. All masks were visually inspected by experimenter blind to cell condition/group. Values obtained using non-fluorescent strains were used as background subtraction.

#### FRET

We followed commonly used methods for quantifying FRET sensor activity^30,56^, and report normalized FRET/CFP ratios. To determine the amount of channel bleed-through between the FRET, CFP, and YFP channels, we imaged cells constitutively expressing only CFP or only YFP in the FRET and CFP channels. This bleed-through value was then subtracted from the FRET/CFP value for each cell.

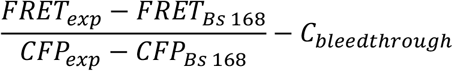

### Spot plating assay

*B. subtilis* strains containing different variants of engineered *lac* repressor in Δ*prpC* and Δ(*prpC-prkC*) backgrounds were grown up in 3 mL LB Lennox to log phase. 5 μL of culture were then spotted onto LB Lennox agar plates and grown overnight. Plates were then imaged in brightfield and mCherry channels on a macroscope device, consisting of a Canon T3i digital single lens reflex (DSLR) camera equipped with a Canon EF-S 60 mm USM lens and controlled by custom-built software. For excitation and emission wavelengths, LEDs with Chroma filters and a Starlight Xpress filter wheel were used, respectively (white light: 3500–4500 K LED; RFP: 588– 592 LED with 572/35 EX and 645/75 EM filter)^57^.

### Bulk fluorescence assays

*B. subtilis* cultures were grown in LB Lennox media inoculated from single colonies. Cultures were grown to late log phase at 37°C in a roller drum, and then diluted 1:30 into fresh LB Lennox in a 96-well plate (150 μL final volume). Plates were grown with continuous shaking at 37°C and measured in a BioTek Synergy NEO or NEO2 plate reader. Measurements of OD_600_ and fluorescence were taken at 10-minute intervals.

#### Calculations

Values obtained from media-only wells and non-fluorescent strains were used as background subtraction for OD_600_ and autofluorescence, respectively. Fluorescence values for each well at each timepoint were normalized to that well’s OD_600_ at each timepoint, yielding RFU/OD_600_ values. For 3 biological replicates the average Δ*prpC* value is then divided by the average Δ(*prpC-prkC*) value to obtain a signal ratio.

### Estimation of PrkC activity through a model of LacI∼P phosphorylation

To model reporter expression as a function of unphosphorylated LacI we use the following Hill function

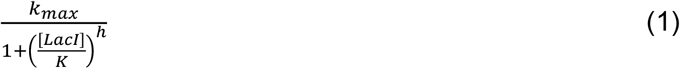

where *k*_*max*_ is the maximum production rate, [*LacI*] denotes the level of unphosphorylated LacI, *h* is the Hill coefficient and parameter *K* represents the level [*LacI*] at which reporter expression is halved. To infer the Hill function, we use data on reporter expression levels for the four different promoter variants synthesizing LacI in Δkinase and Δphosphotase backgrounds. We assume that in the Δkinase all the LacI molecules are in the unphosphorylated state, while in Δphosphotase only a fraction *f* of the total LacI is unphosphorylated. Fitting equation (1) to data estimates the four parameters *(k*_*max*_, *h, K, r*) and reveals a Hill coefficient of *h* ≈ 2.7 and *f* ≈ 0.57 suggesting phosphatase activity even in the Δphosphotase background. The nonlinearity in the Hill function captures the fold change in reporter levels between Δphosphotase and Δkinase with increasing LacI expression (Figure 3E).

Next, we focus on estimating the increase in kinase activity in the presence of antibiotics using the Δphosphotase. Note that the fraction *f* of total LacI that is unphosphorylated is given by

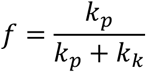

where *k*_*k*_ and *k*_*p*_ are the kinase and phosphatase activity rates, respectively. The kinase activity can be written in terms of *f* as

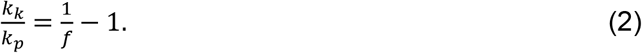

Recall that we estimate *f* ≈ 0.57 in the absence of antibiotics. At 0.5 ug/ml cefotaxime, the increase YFP in levels (normalized by the constitutively expressed CFP) is around 47%±4%where the ± denoted the 95% confidence interval as estimated using bootstrapping. Using the inferred Hill function (1) we back calculate the decrease in unphosphorylated LacI levels needed for an 47% increase in reporter levels. Our analysis shows that the fraction of unphosphorylated LacI decreases from 0.57 (without cefotaxime) to 0.44 (with 0.5 ug/ml cefotaxime) which using equation (2) implies the following increase kinase activity

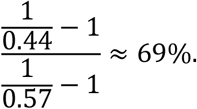

Using bootstrapping we estimate the 95% confidence interval for the increase in kinase activity to be 63% -75%.

## Acknowledgements

This research is based upon work supported in part by the Office of the Director of National Intelligence (ODNI), Intelligence Advanced Research Projects Activity (IARPA) under Finding Engineering Linked Indicators (FELIX) program contract #N660011824505. The views and conclusions contained herein are those of the authors and should not be interpreted as necessarily representing the official policies, either expressed or implied, of ODNI, IARPA, or the U.S. Government. The U.S. Government is authorized to reproduce and distribute reprints for governmental purposes notwithstanding any copyright annotation therein. Additionally, AS acknowledges support from ARO (W911NF1910243) and NIH (R01GM124446). We thank Isaac Plant for the cloning of the LacI∼P variants and FRET sensors as well as Fig 3B. We also thank Jonathan Dworkin for critical reading of the manuscript.

## Data availability

All materials and constructs used in this study are available upon reasonable request from the corresponding author.

## Author Contributions

EAL and CZ performed experiments and data analysis. AL developed a processing pipeline for microscopy images. AS contributed a theoretical model and associated analysis. PAS and EAL designed and coordinated the study. EAL wrote the paper.

