## Supplemental Tables 1-3 for "Protein sensors of bacterial kinase activity reveal antibiotic-dependent kinase activation in single cells"

(Zheng et al, 2021)

**Table S2: Strains Used in this Study**

| Strain | Genotype | Construction | Source |
|---|---|---|---|
| ***B. subtilis* strains** | | | |
| 168 *trpC2* (PB2) | 168 *trpC2* (WT) | | Lab stock & [1] |
| PB702 | PB2 Δ*prpC* | | " |
| PB722 | PB2 Δ(*prpC-prkC*) | | " |
| IP_600 | ΔprpC, sacA::Pveg(cfp-fha2-igaddyvtkpfstr-yfp-stop) CmR | Transformation of IP_325 into PB702 | This study |
| IP_601 | ΔprpC, ΔprkC, sacA::Pveg(cfp-fha2-igaddyvtkpfstr-yfp-stop) CmR | Transformation of IP_325 into PB722 | " |
| IP_358 | ΔprpC, sacA::Pveg(cfp-fha2-igaddyvtkpistr-yfp-stop) CmR | Transformation of IP_326 into PB702 | " |
| IP_362 | ΔprpC, ΔprkC, sacA::Pveg(cfp-fha2-igaddyvtkpistr-yfp-stop) CmR | Transformation of IP_326 into PB722 | " |
| IP_356 | ΔprpC, sacA::Pveg(cfp-fha2-iqedeemtkaipii-yfp-stop) CmR | Transformation of IP_323 into PB702 | " |
| IP_357 | ΔprpC, sacA::Pveg(cfp-fha2-iqedeemakaipii-yfp-stop) CmR | Transformation of IP_319 into PB702 | " |
| IP_360 | ΔprpC, ΔprkC, sacA::Pveg(cfp-fha2-iqedeemtkaipii-yfp-stop) CmR | Transformation of IP_323 into PB722 | " |
| IP_361 | ΔprpC, ΔprkC, sacA::Pveg(cfp-fha2-iqedeemakaipii-yfp-stop) CmR | Transformation of IP_319 into PB722 | " |
| CZ_319 | ΔprpC, sacA::P3(cfp-fha2-iqedeemakaipii-yfp-stop) CmR | Transformation of CZ_159 into PB702 | " |
| CZ_320 | ΔprpC, ΔprkC, sacA::P3(cfp-fha2-iqedeemakaipii-yfp-stop) CmR | Transformation of CZ_159 into PB722 | " |
| CZ_327 | ΔprpC, sacA::P3(cfp-fha2-iqedeemtkaipii-yfp-stop) CmR | Transformation of CZ_163 into PB702 | " |
| CZ_328 | ΔprpC, ΔprkC, sacA::P3(cfp-fha2-iqedeemtkaipii-yfp-stop) CmR | Transformation of CZ_163 into PB722 | " |
| IP_453 | ΔprpC, sacA::Pveg(LacI variant 1) CmR, amyE::Phyperspank(mCherry) SpecR | Transformation of IP_411 and IP_376 into PB702 | " |
| IP_454 | ΔprpC, ΔprkC, sacA::Pveg(LacI variant 1) CmR, amyE::Phyperspank(mCherry) SpecR | Transformation of IP_411 and IP_376 into PB722 | " |
| IP_455 | ΔprpC, sacA::Pveg(LacI variant 2) CmR, amyE::Phyperspank(mCherry) SpecR | Transformation of IP_412 and IP_376 into PB702 | " |
| IP_456 | ΔprpC, ΔprkC, sacA::Pveg(LacI variant 2) CmR, amyE::Phyperspank(mCherry) SpecR | Transformation of IP_412 and IP_376 into PB722 | " |
| IP_457 | ΔprpC, sacA::Pveg(LacI variant 3) CmR, amyE::Phyperspank(mCherry) SpecR | Transformation of IP_413 and IP_376 into PB702 | " |

| | | | |
|---|---|---|---|
| IP_458 | ΔprpC, ΔprkC, sacA::Pveg(LacI variant 3) CmR, amyE::Phyperspank(mCherry) SpecR | Transformation of IP_413 and IP_376 into PB722 | " |
| IP_459 | ΔprpC, sacA::Pveg(LacI variant 4) CmR, amyE::Phyperspank(mCherry) SpecR | Transformation of IP_414 and IP_376 into PB702 | " |
| IP_460 | ΔprpC, ΔprkC, sacA::Pveg(LacI variant 4) CmR, amyE::Phyperspank(mCherry) SpecR | Transformation of IP_414 and IP_376 into PB722 | " |
| IP_461 | ΔprpC, sacA::Pveg(LacI variant 5) CmR, amyE::Phyperspank(mCherry) SpecR | Transformation of IP_415 and IP_376 into PB702 | " |
| IP_462 | ΔprpC, ΔprkC, sacA::Pveg(LacI variant 5) CmR, amyE::Phyperspank(mCherry) SpecR | Transformation of IP_415 and IP_376 into PB722 | " |
| IP_473 | ΔprpC, sacA::Pveg(LacI variant 12) CmR, amyE::Phyperspank(mCherry) SpecR | Transformation of IP_422 and IP_376 into PB702 | " |
| IP_474 | ΔprpC, ΔprkC, sacA::Pveg(LacI variant 12) CmR, amyE::Phyperspank(mCherry) SpecR | Transformation of IP_422, IP_376 into PB722 | " |
| IP_475 | ΔprpC, sacA::Pveg(LacI variant 13) CmR, amyE::Phyperspank(mCherry) SpecR | Transformation of IP_423 and IP_376 into PB702 | " |
| IP_476 | ΔprpC, ΔprkC, sacA::Pveg(LacI variant 13) CmR, amyE::Phyperspank(mCherry) SpecR | Transformation of IP_423, IP_376 into PB722 | " |
| IP_485 | ΔprpC, sacA::Pveg(LacI variant 17) CmR, amyE::Phyperspank(mCherry) SpecR | Transformation of IP_427 and IP_376 into PB702 | " |
| IP_486 | ΔprpC, ΔprkC, sacA::Pveg(LacI variant 17) CmR, amyE::Phyperspank(mCherry) SpecR | Transformation of IP_427, IP_376 into PB722 | " |
| CZ_12 | ΔprpC, sacA::P1(lacI-P) Phyperspank(YFP) CmR | Transformation of CZ_8 into PB702 | " |
| CZ_16 | ΔprpC, ΔprkC, sacA::P1(lacI-P) Phyperspank(YFP) CmR | Transformation of CZ_8 into PB722 | " |
| CZ_13 | ΔprpC, sacA::P2(lacI-P) Phyperspank(YFP) CmR | Transformation of CZ_9 into PB702 | " |
| CZ_14 | ΔprpC, sacA::P3(lacI-P) Phyperspank(YFP) CmR | Transformation of CZ_10 into PB702 | " |
| CZ_15 | ΔprpC, sacA::P4(lacI-P) Phyperspank(YFP) CmR | Transformation of CZ_11 into PB702 | " |
| CZ_17 | ΔprpC, ΔprkC, sacA::P2(lacI-P) Phyperspank(YFP) CmR | Transformation of CZ_9 into PB722 | " |
| CZ_18 | ΔprpC, ΔprkC, sacA::P3(lacI-P) Phyperspank(YFP) CmR | Transformation of CZ_10 into PB722 | " |
| CZ_19 | ΔprpC, ΔprkC, sacA::P4(lacI-P) Phyperspank(YFP) CmR | Transformation of CZ_11 into PB722 | " |

| | | | |
|---|---|---|---|
| IP_517 | ΔprpC, sacA::Ppcn(lacI-P) Phyperspank(YFP) CmR | Transformation of IP_502 into PB702 | " |
| IP_518 | ΔprpC, ΔprkC, sacA::Ppcn(lacI-P) Phyperspank(YFP) CmR | Transformation of IP_502 into PB722 | " |
| CZ678 | sacA::P1(lacI-P) CmR, amyE:: Pveg(CFP) KanR | Transformation of JN_11 and CZ_8 into Bs. 168 | " |
| CZ679 | ΔprpC, sacA::P1(lacI-P) CmR, amyE:: Pveg(CFP) KanR | Transformation of JN_11 and CZ_8 into PB702 | " |
| CZ680 | ΔprpC, ΔprkC, sacA::P1(lacI-P) CmR, amyE:: Pveg(CFP) KanR | Transformation of JN_11 and CZ_8 into into PB722 | " |

**Table S2: Plasmids Used in this Study**

| Plasmid | Genotype | Construction | Source |
|---|---|---|---|
| pSac-cm | Shuttle vector for integration at *sacA* | | Lab stock[2] |
| CZ_8 | P1(LacI-P) Phyperspank(YFP) Cm, for integration into sacA locus | Assembled via Golden Gate cloning. Promoter added as annealed oligos with complementary overhangs (IP_P_1296, 1297, 1298, 1299). Backbone amplified with IP_P_1300, 1293 from IP_502. | This work |
| CZ_9 | P2(LacI-P) Phyperspank(YFP) Cm, for integration into sacA locus | Assembled via Golden Gate cloning. Promoter added as annealed oligos with complementary overhangs (IP_P_1296, 1297, 1301, 1302). Backbone amplified with IP_P_1300, 1293 from IP_502. | This work |
| CZ_10 | P3(LacI-P) Phyperspank(YFP) Cm, for integration into sacA locus | Assembled via Golden Gate cloning. Promoter added as annealed oligos with complementary overhangs (IP_P_1296, 1297, 1303, 1304). Backbone amplified with IP_P_1300, 1293 from IP_502. | This work |
| CZ_11 | P4(LacI-P) Phyperspank(YFP) Cm, for integration into sacA locus | Assembled via Golden Gate cloning. Promoter added as annealed oligos with complementary overhangs (IP_P_1296, 1297, 1305, 1306). Backbone amplified with IP_P_1300, 1293 from IP_502. | This work |

| | | | |
|---|---|---|---|
| CZ_159 | Pveg7(cfp-fha2-iqedeemakaipii-yfp-stop) Cm, for integration into sacA locus | Assembled via Golden Gate cloning. Promoter added as annealed oligos with complementary overhangs (IP_P_1296, 1297, 1303, 1304). Backbone amplified with CZp148, CZp149 from IP_319. | This work |
| CZ_163 | Pveg7(cfp-fha2-iqedeemtkaipii-yfp-stop) Cm, for integration into sacA locus | Assembled via Golden Gate cloning. Promoter added as annealed oligos with complementary overhangs (IP_P_1296, 1297, 1303, 1304). Backbone amplified with CZp148, CZp149 from IP_323. | This work |
| IP_319 | Pveg(cfp-fha2-iqedeemakaipii-yfp-stop) CmR, for integration at sacA locus | direct synthesis into pSac-cm | This work |
| IP_323 | Pveg(cfp-fha2-iqedeemtkaipii-yfp-stop) CmR, for integration at sacA locus | " | This work |
| IP_324 | Pveg(cfp-fha2-stop-yfp-stop) cm, for integration at sacA locus | " | This work |
| IP_325 | Pveg(cfp-fha2-igaddyvtkpfstr-yfp-stop) CmR, for integration at sacA locus | " | This work |
| IP_326 | Pveg(cfp-fha2-igaddyvtkpistr-yfp-stop) CmR, for integration at sacA locus | " | This work |
| IP_376 | Phyperspank(mCherry) SpecR, for integration into amyE | Golden gate cloning described in detail[3] and vector maps available at http://nrs.harvard.edu/urn-3:HUL.InstRepos:42029477 | This work |
| IP_411 | Pveg(LacI variant 1) CmR, for insertion at sacA locus | " | This work |
| IP_412 | Pveg(LacI variant 2) CmR, for insertion at sacA locus | " | This work |
| IP_413 | Pveg(LacI variant 3) CmR, for insertion at sacA locus | " | This work |

| | | | |
|---|---|---|---|
| IP_414 | Pveg(LacI variant 4) CmR, for insertion at sacA locus | “ | This work |
| IP_415 | Pveg(LacI variant 5) CmR, for insertion at sacA locus | “ | This work |
| IP_422 | Pveg(LacI variant 12) CmR, for insertion at sacA locus | “ | This work |
| IP_423 | Pveg(LacI variant 13) CmR, for insertion at sacA locus | “ | This work |
| IP_427 | Pveg(LacI variant 17) CmR, for insertion at sacA locus | “ | This work |
| IP_502 | Ppcn(LacI-P) Phyperspank(YFP) Cm, for integration into sacA locus | “ | This work |
| JN_11 | Pveg(CFP) KanR, for integration at amyE locus | Assembled via Golden Gate cloning. CFP insert amplified with JNp_23, JNp_24. Backbone amplified from IP_193 with JNp_1, JNp_2. | This work |

**Table S3: Oligos used in this study**

| Name | Sequence (5'-3') |
|---|---|
| IP_P_1296 | ACCCAATTTTGTCAAAATAATTTTATTGACAACGTCTTATTAACGTTGATACCGGTTAA |
| IP_P_1297 | AAATTTAACCGGTATCAACGTTAATAAGACGTTGTCAATAAAATTATTTTGACAAAATT |
| IP_P_1298 | ATTTTATTTGACAAAAATGGGCTCGTGTTGGACAATAAATGTGGAGAAAAGCTAGCGATT |
| IP_P_1299 | AGTTAATCGCTAGCTTTTCTCCACATTTATTGTCCAACACGAGCCCATTTTTGTCAAATA |
| IP_P_1301 | ATTTTATTTGACAAAAATGGGCTCGTGTTGTTGAATAAATGTGGAGAAAAGCTAGCGATT |
| IP_P_1302 | AGTTAATCGCTAGCTTTTCTCCACATTTATTCAACAACACGAGCCCATTTTTGTCAAATA |
| IP_P_1303 | ATTTTATTTGACAAAAATGGGCTCGTGTTGAACAATAAATGTGGAGAAAAGCTAGCGATT |
| IP_P_1304 | AGTTAATCGCTAGCTTTTCTCCACATTTATTGTTCAACACGAGCCCATTTTTGTCAAATA |
| IP_P_1305 | ATTTTATTTGACAAAAATGGGCTCGTGTTGTCCAATAAATGTGGAGAAAAGCTAGCGATT |
| IP_P_1306 | AGTTAATCGCTAGCTTTTCTCCACATTTATTGGACAACACGAGCCCATTTTTGTCAAATA |
| CZp148 | CATGTCGGTCTCCGGGTATCCGACCATTGACTGCC |
| CZp149 | CATGTCGGTCTCCAACTAATAAGGAGGACAAACATG |
| JNp_1 | CGTCTCTCTGCCTCCTCATCCTCTTC |
| JNp_2 | CGTCTCGGCTCCGTCGATACTATGTTA |
| JNp_23 | CGTCTCTGCAGAATTTTGTCAAAATAATTTTATTGACAA |
| JNp_24 | CGTCTCGGAGCTTAAGTGCCCTTATACAACTCG |
